# A cell-to-patient machine learning transfer approach uncovers novel basal-like breast cancer prognostic markers amongst alternative splice variants

**DOI:** 10.1101/2020.11.12.380485

**Authors:** Jean-Philippe Villemin, Claudio Lorenzi, Andrew Oldfield, Marie-Sarah Cabrillac, William Ritchie, Reini F. Luco

## Abstract

**Background:** Breast cancer is amongst the 10 first causes of death in women worldwide. Around 20% of patients are misdiagnosed leading to early metastasis, resistance to treatment and relapse. Many clinical and gene expression profiles have been successfully used to classify breast tumours into 5 major types with different prognosis and sensitivity to specific treatments. Unfortunately, these profiles have failed to subclassify breast tumours into more subtypes to improve diagnostics and survival rate. Alternative splicing is emerging as a new source of highly specific biomarkers to classify tumours in different grades. Taking advantage of extensive public transcriptomics datasets in breast cancer cell lines (CCLE) and breast cancer tumours (TCGA), we have addressed the capacity of alternative splice variants to subclassify highly aggressive breast cancers.

**Results:** Transcriptomics analysis of alternative splicing events between luminal, basal A and basal B breast cancer cell lines identified a unique splicing signature for a subtype of tumours, the basal B, whose classification is not in use in the clinic yet. Basal B cell lines, in contrast with luminal and basal A, are highly metastatic and express epithelial-to-mesenchymal (EMT) markers, which are hallmarks of cell invasion and resistance to drugs. By developing a semi-supervised machine learning approach, we transferred the molecular knowledge gained from these cell lines into patients to subclassify basal-like triple negative tumours into basal A- and basal B-like categories. Changes in splicing of 25 alternative exons, intimately related to EMT and cell invasion such as ENAH, CD44 and CTNND1, were sufficient to identify the basal-like patients with the worst prognosis. Moreover, patients expressing this basal B-specific splicing signature also expressed newly identified biomarkers of metastasis-initiating cells, like CD36, supporting a more invasive phenotype for this basal B-like breast cancer subtype.

**Conclusions:** Using a novel machine learning approach, we have identified an EMT-related splicing signature capable of subclassifying the most aggressive type of breast cancer, which are basal-like triple negative tumours. This proof-of-concept demonstrates that the biological knowledge acquired from cell lines can be transferred to patients data for further clinical investigation. More studies, particularly in 3D culture and organoids, will increase the accuracy of this transfer of knowledge, which will open new perspectives into the development of novel therapeutic strategies and the further identification of specific biomarkers for drug resistance and cancer relapse.

## BACKGROUND

Breast cancer is a heterogenous disease with multiple molecular drivers and disrupted regulatory pathways that impact the therapeutic strategy to choose for an optimal treatment (1,2). The development of large-scale genomics and transcriptomics methods has increased the capacity to identify clinically-relevant tumour subtypes with distinct molecular signatures. These can be used for a better choice of treatment and/or prediction of potential metastasis which can improve survival outcome (3,4). However, patients are still facing a high percentage of misdiagnosis in which undetected early metastasis and/or inappropriate choice of treatment can lead to deadly complications by the use of unnecessary severe chemotherapies or the generation of drug resistance and tumour relapse (5). Currently, breast cancer is classified into four major categories (luminal A, luminal B, Her2-positive and basal-like) based on expression of three receptors: oestrogen and progesterone hormonal receptors (ER and PR) and the epidermal growth factor receptor ERBB2 (Her2). The most aggressive tumours are basal-like. They are mostly negative for the three receptors, and thus called triple negative breast cancer (TNBC), and represent 10−20% of all breast cancers. These tumours are usually found in younger patients with a larger size and higher probability of lymph node infiltration and metastasis (2,6). Furthermore, the absence of all three receptors reduces the number of targeted therapeutic strategies to be used, leaving nonspecific chemotherapy as the standard treatment of choice, which soon leads to dose-limiting side-effects, resistance to treatment and finally clinical relapse in less than 5 years (6). A better understanding of the molecular differences between these tumour categories is thus necessary to improve treatment choice and the detection of early metastasis and tumour relapse, which will significantly impact patient’s outcome.

There have been many attempts to identify novel therapeutic targets and/or prognostic biomarkers to better subclassify breast cancer tumours (7). Over 170 independent breast cancer susceptibility genomic variants have been identified. Many of which have been associated with a specific tumour category, such as ER positiveness or Her2 amplification. However no clear subcategories exist despite tumour heterogeneity and differences in clinical response to treatment and tumour relapse within the same category (8–10). Interestingly, alternative splicing is an emerging source of new biomarkers and therapeutic targets in cancer (11–15).

The alternative processing of mRNA precursors enables one gene to produce multiple protein isoforms with different functions, increasing protein diversity and the capacity of a cell to adapt to new environments. Cancer cells take advantage of this mechanism to produce new proteins with added, deleted, or altered functional domains that confer a selective advantage for tumour proliferation (16), resistance to apoptosis (17), stimulation of angiogenesis (18), metabolic adaptation (19,20) and acquisition of migratory and invasive phenotypes for distal metastasis (13,21–24). The existence of functionally relevant cancer specific protein isoforms is therefore a promising new source of highly specific and less toxic therapeutic targets for the development of isoform-specific antibodies and/or splice-switching antisense oligonucleotides (25,26).

By taking advantage of an extensive transcriptomics and anti-tumour compound screening information publicly available in cancer cell lines from the Cancer Cell Line Encyclopedia (CCLE) (27), we identified a splicing signature that can stratify basal breast cancer cell lines into two well-known subtypes, basal A and basal B. This splicing signature is unique to basal B cells, which are the most aggressive type of cells with a highly invasive and stem-like phenotype. Interestingly this basal B-specific splicing signature is also characteristic of a biological process called the epithelial-to-mesenchymal transition (EMT), in which epithelial cells acquire mesenchymal features that are advantageous for the cancer cell, such as increased cell motility to invade distal organs in metastasis, resistance to apoptosis, refractory responses to chemotherapy and immunotherapy, and acquisition of stem cell-like properties like in cancer stem cells (28,29). Using a semi-supervised machine learning classifier, we subclassified basal-like patients from the Cancer Genome Atlas (TCGA) (30) into two groups based on expression of this basal B-specific splicing signature. 45% of basal-like breast cancer patients (85/188) shared a unique 25 spliced gene signature characteristic of EMT. In this signature, we found well-known markers of malignancy, such as ENAH EMT splice variant that promotes lung metastasis (31) or CSF1 variant which promotes macrophage infiltration and distal metastasis (32), together with new promising splicing candidates of tumour progression and invasiveness. Finally, expression of this basal B signature was sufficient to identify the triple negative breast cancer tumours with poor survival, highlighting the prognostic value of the newly identified splicing biomarkers to subclassify one of the most heterogenous and difficult to treat type of breast cancer. More studies in cell lines, particularly regarding resistance to treatment and cell invasion will be essential to refine this splicing signature in view of orienting treatment or predicting metastasis sites.

In conclusion, by adapting a machine learning approach, we were able to transfer the molecular knowledge obtained in experimental cell lines to identify novel biomarkers of poor prognosis and metastasis amongst triple negative breast cancers in patients. Furthermore, the study of the regulatory pathway responsible for this specific splicing signature pointed to RBM47 as a major regulator for which differential expression levels also correlate with distinct prognostic values, turning this splicing regulator into a promising novel therapeutic target. Further clinical and functional validation of the 25 splicing events proposed in our basal B-specific splicing signature will open new perspectives in the understanding of triple negative breast cancers and the improvement of currently available therapeutic strategies and survival outcome.

## RESULTS

### A distinctive Basal B-like breast cancer splicing signature

Data mining of large-scale genomics and transcriptomics datasets in breast cancer cell lines are a promising source of novel biomarker and therapeutic targets (23,33,34). We sought to leverage the wealth of transcriptomics and functional data available in cancer cell lines to better understand different profiles of breast cancer. Hierarchical clustering of changes in alternative splicing of cassette exons and gene expression profile of 80 breast cancer cell lines from two extensive and complementary projects (Table S1) revealed basal B cell lines as a distinctive group of cells with an expression and splicing profile significantly different from basal A and luminal cancer cells (Fig.1). In contrast to basal-like breast cancer patients, basal breast cancer cell lines are divided into two subgroups, basal A and basal B, depending on the expression profile of a subset of basal (cytokeratins, integrins), stem cell (CD44, CD24) and mesenchymal markers (Vimentin, fibronectin, MSN, TGFBR2, collagens, proteases) (35). Basal B cell lines are mostly triple negative breast cancer cells that express classical mesenchymal and stem cell markers characteristic of the epithelial-to-mesenchymal transition (EMT), a biological process intimately related to cancer progression, resistance to treatment and metastasis (28,35). In concordance, basal B cells are morphologically less differentiated, with a mesenchymal-like shape, and a more invasive phenotype in culture assays than basal A and luminal cells (35–37). To identify the transcriptional signature characteristic of basal B cells, we repeated the hierarchical clustering in just basal A and basal B cell lines in the two projects to merge all the differentially expressed and spliced transcripts responsible for the segregation of basal B cell lines (Fig.S1). We found 635 genes and 217 spliced isoforms with significantly different levels between basal A and basal B cells (Fig.S1a,b). Remarkably, when assessing the gene overlap between differentially expressed and differentially spliced genes, we found that differentially spliced genes were rarely affected at the transcriptional level, suggesting that two different subsets of genes, and thus regulatory layers, are responsible for the basal B-like phenotype (Fig.S1c).

**Figure 1.**
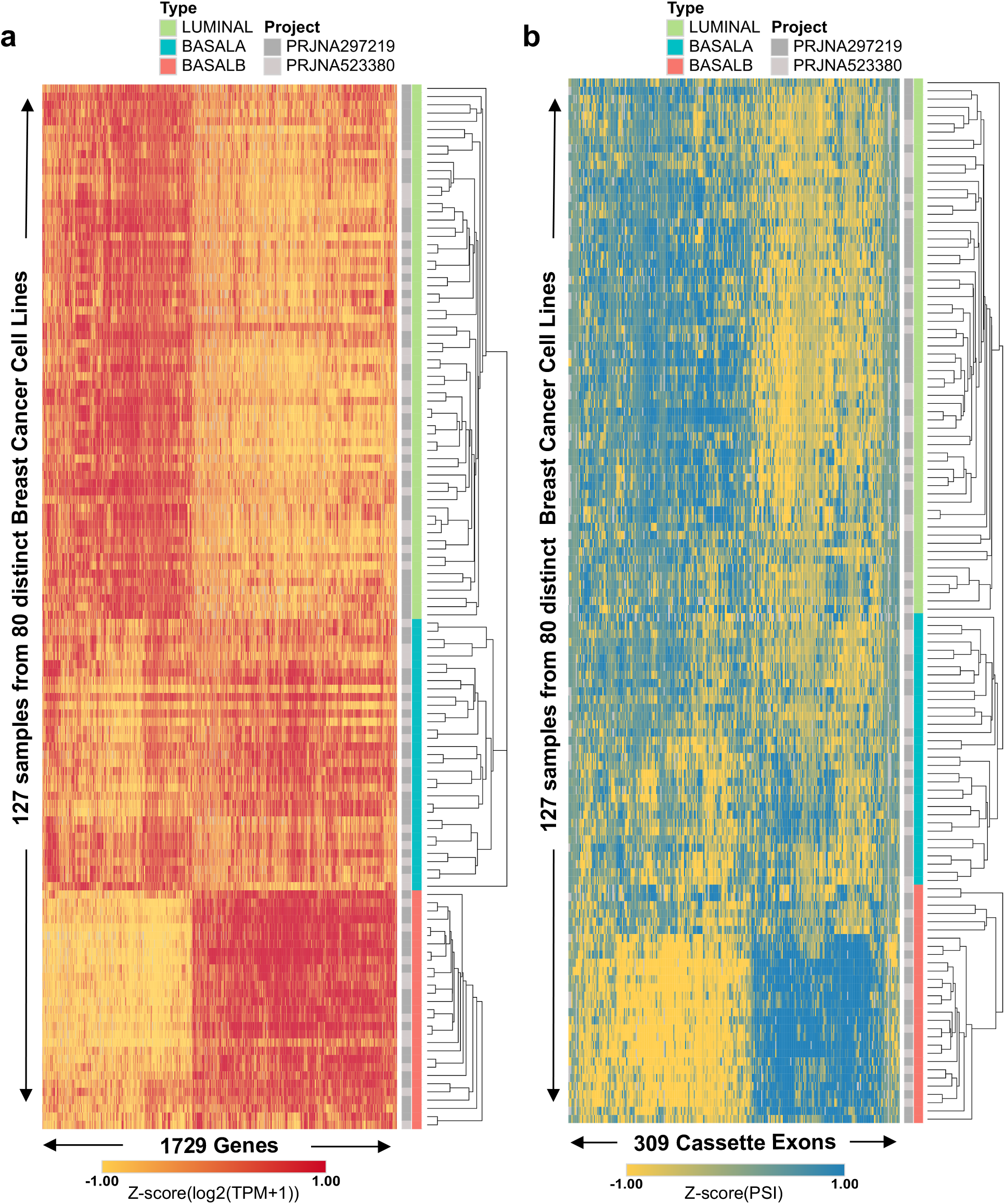
Differential clustering of basal B cell lines based on gene expression and splicing patterns. **a.** Heatmap of gene expression levels, in Transcripts per Million (TPM) values, of 1729 genes differentially regulated between Luminal, basal A and basal B cell lines (P-value < 10^−5^ by Kruskal-Wallis Test). **b.** Heatmap of exon inclusion levels, using Percentage Spliced-In (PSI), of 309 exons differentially spliced between luminal, basal A and basal B cell lines (P-value <10^−5^ by Kruskal-Wallis Test).

Gene set enrichment analysis (GSEA) (38) between basal B and basal A cells confirmed the stem cell-like EMT phenotype characteristic of basal B cell lines (Fig.2a,b), which was supported with a higher CD44+/CD24− stem cell score (Fig.2e). In parallel, gene ontology analysis, using DAVID (39), of differentially spliced genes also underlined biological terms that are hallmarks of EMT and cell invasiveness, such as GTPase activity, anchoring and adherens junction, cadherin binding involved in cell-cell adhesion and cytoskeletal protein binding (Fig.2a,d). Importantly, another malignant characteristic acquired by cancer cells undergoing EMT is resistance to chemotherapy, which often leads to clinical relapse. Gene set enrichment analysis found upregulation of genes resistant to the Epidermal Growth Factor Receptor (EGFR) inhibitor Gefitinib (Fig.2c), which is an alternative to hormonal therapy in Her2+ breast cancer tumours, but does not seem as efficient in triple negative tumours, such as basal B-like (40). Available drug assays from the Genome Drug Sensitivity in Cancer portal (GDSC) (41) confirmed the need of a higher concentration (IC50) of Gefitinib, and other EGFR inhibitors (Erlotinib, Sapitinib), to have the same deleterious effect on basal B compared to basal A cancer cells (Fig.2f). Basal B cell lines also showed a significant resistance to well-known inhibitors of the cell cycle (Irinotecan, Taselisib, 5-Fluorouracil), drug inducers of cell death (AZD5582, AZD5991) and other receptor tyrosine kinase inhibitors, such as Savolitinib that inhibits c-MET to reduce tumour persistence and metastasis (42).

**Figure 2.**
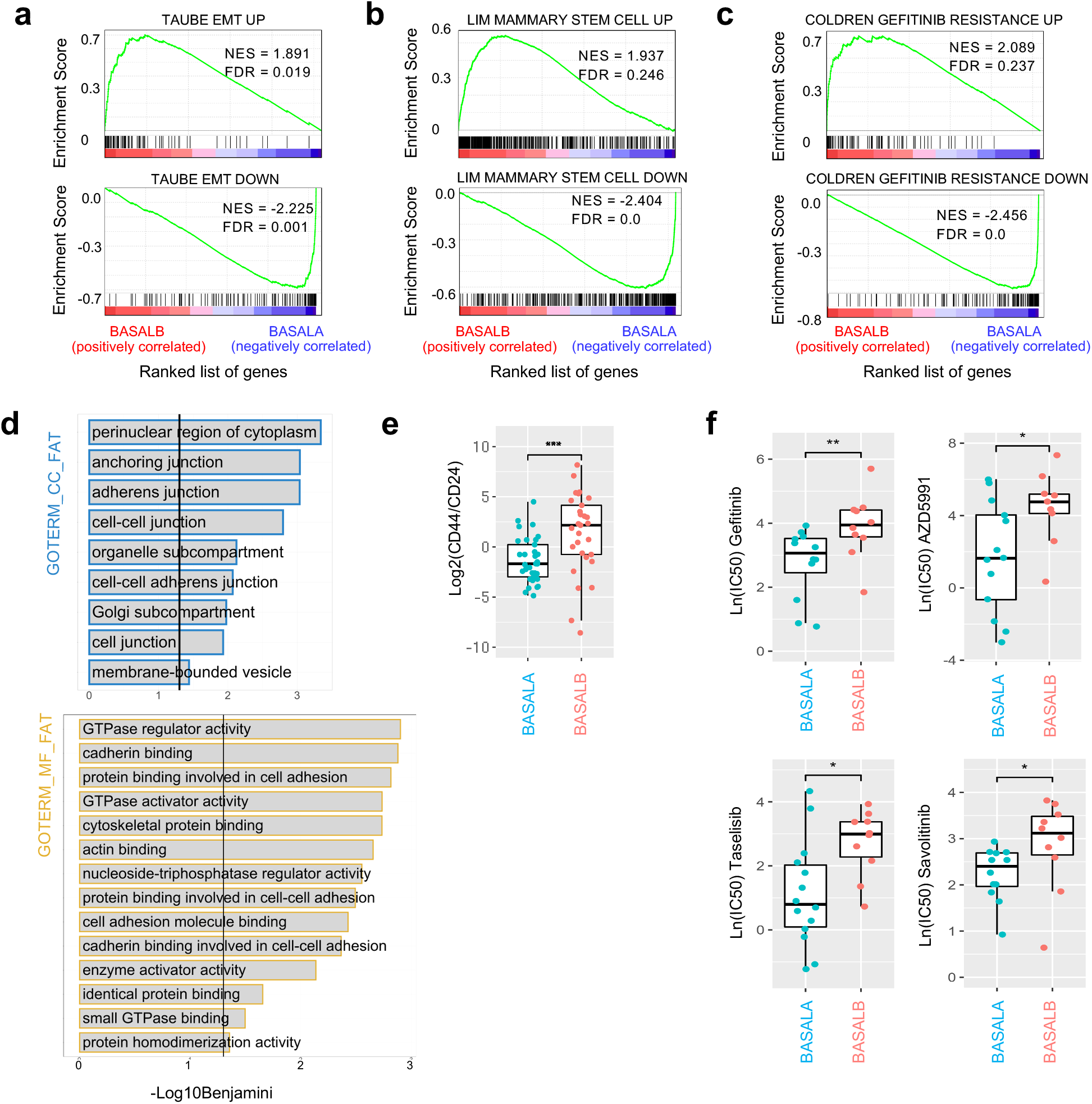
Basal B cell lines show mesenchymal, stem-like and resistance to treatment characteristics. **a,b,c.** Gene Set Enrichment Analysis (GSEA) of differentially expressed genes between basal A and B cell lines for three different signatures: Mammary Stem Cell, EMT and Resistance to Gefitinib. Up-regulated genes in all signatures are enriched in basal B cell lines (FDR<0.25). **d**. Gene ontology analysis bar graphs for differentially spliced genes between basal A and B cell lines. Gene ontology terms related to Cellular Component (GO_CC_FAT) and Molecular Function (GO_MF_FAT) are shown in the y axis in blue and yellow, respectively. Benjamini false discovery rate (FDR, −log10) is shown on the x axis. Vertical lines mark an FDR=0.05. **d.** Box plots of the median and 25th percentile of the CD44/CD24 log2 expression ratio for basal A and B cell lines. P-value is calculated using the Wilcoxon rank-sum test. **f.** Boxplots comparing IC50 values in basal A and B cell lines upon treatment with different drugs from the Genomics of Drug Sensitivity in Cancer 2 (GDS2) dataset. P-values are calculated using the Wilcoxon rank-sum test.

In summary, we have identified two distinct transcriptional and splicing signatures, specific of basal B cell lines, that underline an EMT phenotype with molecular characteristics related to cell invasion, stemness and resistance to chemotherapy. We next sought to investigate whether this basal B-specific splicing signature could also be used to subclassify basal-like/triple negative breast cancer patients.

### A semi-supervised machine learning approach to subclassify basal-like breast cancer patients

As a first and simple approach, we performed a hierarchical clustering followed by a k-means clustering (k=2 for “A-like” and “B-like”) of the 188 patients, annotated as basal-like in The Cancer Genome Atlas Program (TCGA), using the 635 differentially expressed or 217 differentially spliced cassette exons characteristic of basal B cell lines (Fig. S2a,b). Using such a method, patients are forced to be classified into one of the two groups based on differences in gene expression or splicing patterns. Since basal B cell lines show more invasive, cancer stem cell-like phenotypes, we assessed whether these aggressive characteristics were translated to the “B-like” patient group. We thus tested for differences in disease specific survival (DSS) rates with the hypothesis that “B-like” patients should have poor prognosis. Kaplan-Meier analysis of DSS did not show significant differences between the two subgroups of basal-like patients (Fig. S2c,d). However, we did observe a tendency for “B-like” patients to have a poor survival compared to “A-like” when just looking at differences in splicing, contrary to expression levels (p-value=0.09 vs 0.57, respectively – Fig. S2c,d).

It was not surprising that the transcript-level and splicing signatures did not translate directly from simplistic cell culture models to much more complex tumour patients with specific cell micro-environments and differences in cell heterogeneity. However, because the patients showed clear “A-like” and “B-like” signatures, we sought to develop a machine learning approach that would allow us to transfer part of the molecular and phenotypic observations found in cell-lines to patient data. Transfer learning is a recent research problem that focuses on storing knowledge gained while solving one problem and applying it to a different but related problem. Because we wanted to ensure that the newly developed cell-to-patient transfer learning algorithm could create interpretable models, we used a decision tree-based approach called Random Forest. In this cell-to-patient random forest classification method, we started by classifying basal A or basal B cell-lines based on their splicing and expression profile (Fig. 3a). Then, once the model was trained on cell-lines, we would start integrating patient data gradually into the model. This was done iteratively by integrating at each round of classification the patients best predicted to be basal A-like and basal B-like, so their added informative value could be used back to train the system and improve the next round of classification (Fig.3a). With this semi-supervised approach, the probability of assigning a patient to a specific subgroup can evolve and improve at each round based on the updated information obtained from the best predicted patients, reaching at the end a stable population with the labels ‘basal A-like’, ‘basal B-like’ or ‘unclassified’ determined by the algorithm after 10^−12^ rounds (Fig.3b,c). Out of the 188 basal-like patients, 85 were classified as basal B-like, 78 as basal A-like and 25 could not be classified based on their splicing signature. Importantly, when applying the same strategy, but using differentially expressed genes as classifying features, only 4 patients were classified as basal B-like and the rest as basal A-like (Fig. S3). Interestingly all the patients sharing a common basal B-specific transcriptional signature were also lowly expressing well-known markers of aggressive breast cancer, which are claudins. Claudin-low breast cancer is a molecular subtype of breast cancer originally identified by gene expression profiling and reportedly associated with higher tumour grade, extensive lymphocytic infiltrate and poor survival (43). Even though claudin-low tumours preferentially display a basal-like EMT phenotype, only a minority of triple-negative breast cancers are actually claudin-low, proving a limited clinical value to identify aggressive triple negative breast cancer tumours (43). The basal B-specific transcriptional signature transferred from breast cancer cell lines reproduced this claudin-low classification (Fig.S3a). However, the newly identified splicing-specific basal B-like signature went beyond this claudin-low classification and identified more patients with a common signature different from claudin-low, supporting a potential new clinical value to subclassify basal-like tumours with a distinct molecular phenotype (Fig.4a).

**Figure 3.**
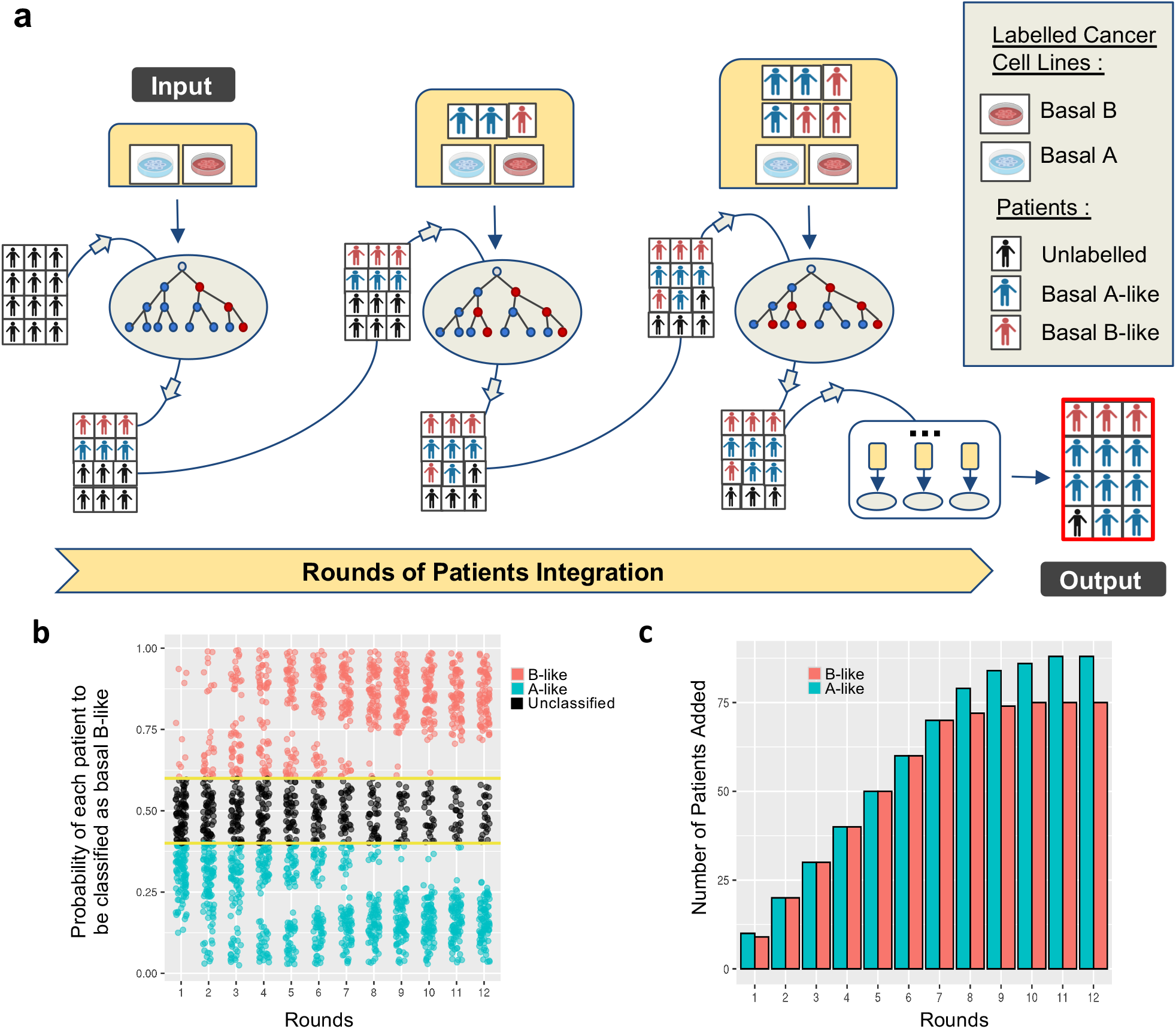
A Random Forest Classifier using knowledge transfer from cell lines to patients. **a.** Workflow scheme: a random forest (RF) model is built using cell lines labelled as Basal B (red) or Basal A (blue). It is then run iteratively, integrating at each round patients whose probability to be classified in one group or the other is amongst the ten highest. The classifier stops when no more patients can be classified. **b.** Probability of a basal-like patient to be classified as basal B-like, basal A-like or unclassified over each round. Yellow lines indicate thresholds used to classify a patient as basal B-like (>0.6) or basal A-like (<0.4) **c.** Bar plot of the number of patients added through each round. At each round, patients with the highest probability to be classified are sequentially incorporated to the input cell lines in order to create a new classifier for the next round of integration.

**Figure 4.**
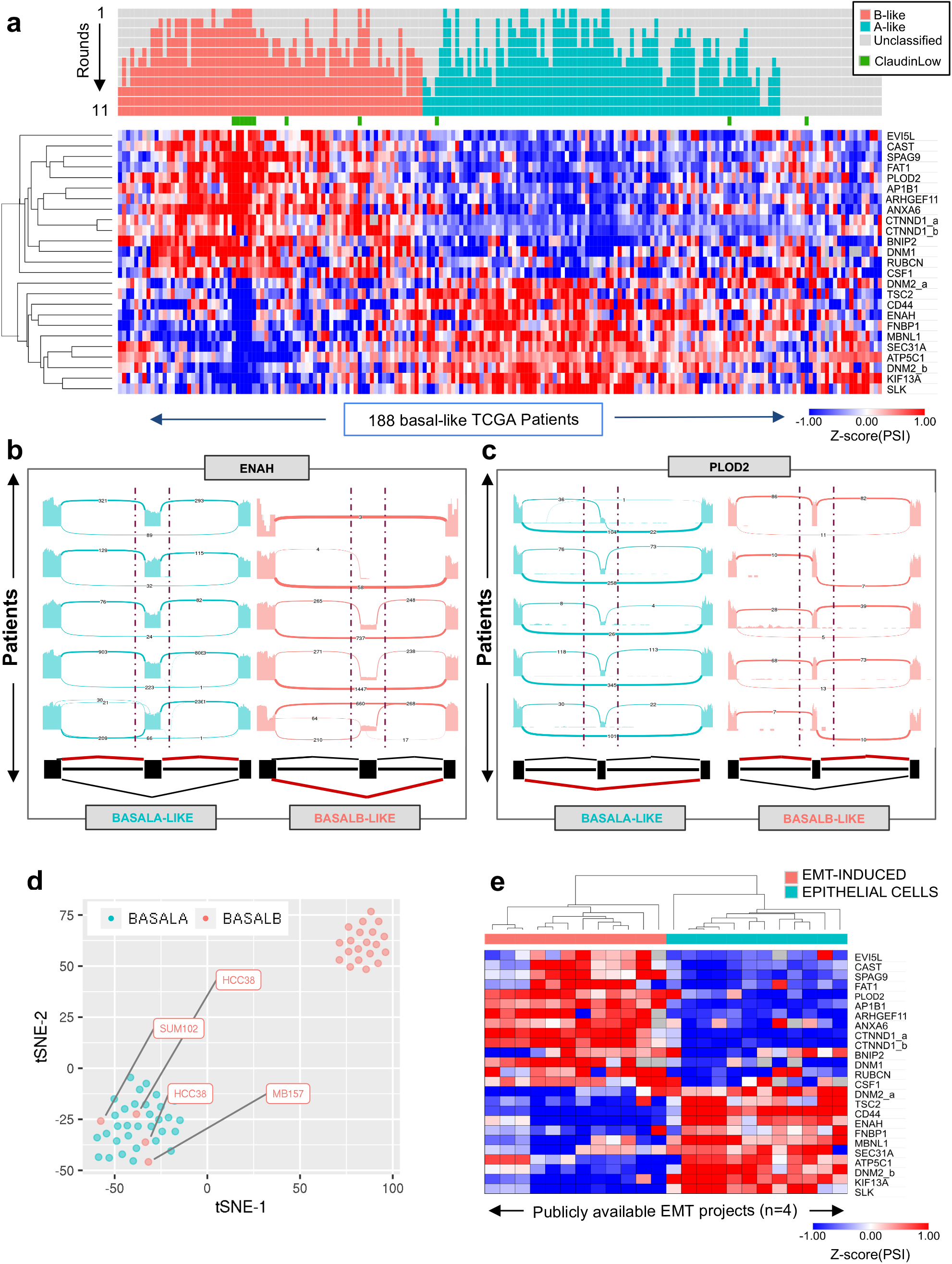
The basal B-specific splicing signature is associated to EMT features. **a.** Heatmap of the Percentage Spliced-In (PSI) values of the 25 cassette exons most informative to classify TCGA basal-like patients into basal B-like (red) or basal A-like (blue). Claudin low tumors are highlighted in green. **b,c.** Sashimi plots displaying ENAH and PLOD2 splicing patterns in randomly selected patients classified as basal A-like and basal B-like. **d.** Changes in alternative splicing of these 25 basal B-specific splicing events is sufficient to properly cluster 55 basal breast cancer cell lines from 3 unrelated sequencing projects into basal B and basal A using t-SNE. **e.** Heatmap of the PSI values of the 25 basal B-specific splicing signature in public RNA-seq datasets from four different EMT projects. Basal B-like events have the same splicing patterns as EMT-induced cells.

### An EMT-related basal B-specific splicing signature that marks poor prognosis

To extract the most informative features from the cell-to-patient transfer learning classifier, we used the Boruta feature selection method (44). This allowed us to select the most informative splicing events responsible for the basal A/B classification (Fig.4a). Out of the 217 differentially spliced exons between basal A/B cell lines, just 25 were needed to subclassify breast cancer patients in basal A or basal B-like tumours (Fig.4a and Table S2). Sashimi plots showing splicing patterns of some of these basal B-specific splicing events in patients, such as the well-known splicing biomarker of cancer metastasis ENAH (26) and the newly identified splicing biomarkers PLOD2, SPAG9 and KIF13a, validated the observed changes in splicing between basal A and basal B-like patients (Fig.4b-c and Fig.S4a-b). Moreover, the changes in percentage of spliced-in (PSI) of the 25 basal B-specific splicing events between the two subtypes of basal-like patients correlated with the observed splicing changes between basal A/B cell lines (Fig.S4c-d), further supporting the transfer of knowledge from the laboratory to the clinic. Finally, distribution of 52 breast cancer cell lines from three independent sequencing projects different from the ones used for the training of the semi-supervised classifier (Table S1), showed a 93% accuracy in the spatial segregation (t-SNE) of basal A from basal B cells based on the splicing pattern of the 25 newly identified splicing events, which validated our methodology and the specificity of the splicing signature towards a basal B-like phenotype (Fig.4d).

Consistent with basal B cell lines being more mesenchymal, differences in the alternative splicing of these 25 basal B-specific splicing events in four different cellular models of EMT, coming from different cell types and methods of EMT induction (45–48), successfully clustered epithelial cells from EMT-induced cells with a pattern of splicing equivalent to basal A and basal B-like patients, respectively and as expected (Fig.4e). Of note, another 25 gene-based EMT-like splicing signature characteristic of luminal breast cancer tumours has also been identified capable of subclassifying mesenchymal-like breast cancer tumours with poor prognosis (33). Consistent with a more luminal-specific signature, despite both marking EMT phenotypes, not more than six splicing events were found in common between the two splicing signatures (ATP5C1, CTNND1, KIF13a, PLOD2, SEC31a and SPAG9), which further supports the specificity of our newly identified splicing signature for basal-like triple negative breast cancer. Since luminal tumours expressing this EMT-like splicing signature had a poor prognosis compared to epithelial ones, we next assessed the disease specific survival rate of basal A-like and basal B-like patients based on expression of the newly identified splicing signature. Consistent with an EMT-like, drug resistant, invasive signature, basal B-like patients had a poor prognosis compared to basal A-like patients (log-rank test p = 0.0067, HR = 4.87; IC95%: [1.37-17.28] in Kaplan-Meier analysis and univariate Cox regression) (Fig.5a). Moreover, using one of the first established molecular subtypes of triple negative breast cancer tumours based on gene expression, which is the Lehman classification (49), we found that basal B-like patients are mostly found in the categories associated with mesenchymal (M), Mesenchymal stem-like (MSL) and Immunomodulatory (IM) subtypes (Fig.5b), which is consistent with a gene set enrichment of terms related to inflammatory responses and hallmark of EMT (Fig.5c).

**Figure 5.**
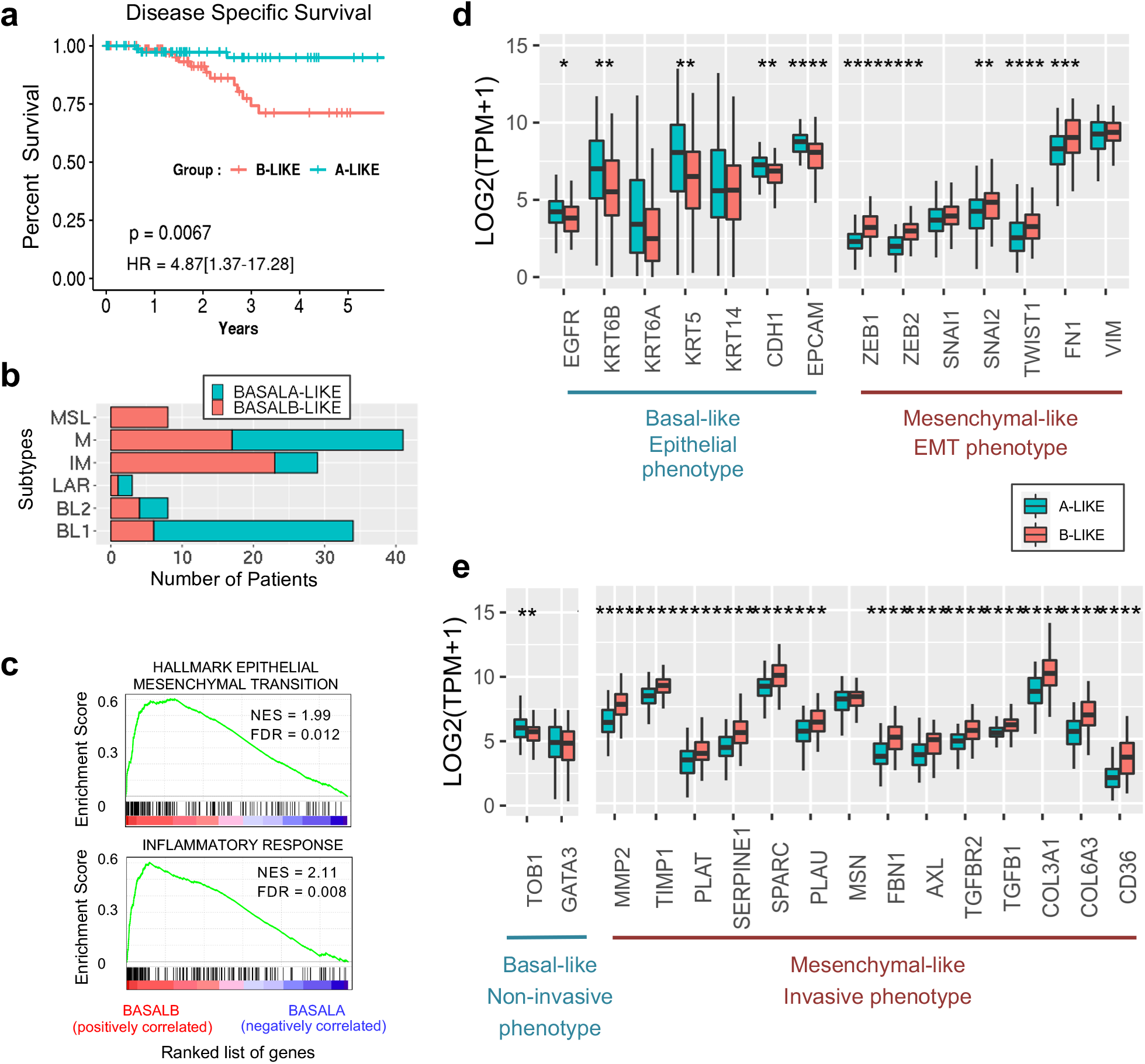
Basal B-like patients express hallmarks of EMT and metastasis that leads to a poor prognosis. **a.** Kaplan-Meier plots of disease specific survival in basal A-like (blue) and basal B-like patients (red). Hazard ratio (HR) and logrank p-value (P) discriminating the two groups are shown. **b.** Lehman classification for basal A- and B-like patients. **c.** Gene Set Enrichment Analysis (GSEA) of the genes differentially expressed between basal A- and B-like patients. Hallmark EMT and inflammatory response signatures are enriched in basal B-like patients. **d.** Box plots of the median and 25th percentile of the expression levels (in TPM) of major epithelial and mesenchymal-like EMT markers in basal A-like (blue) and basal B-like (red) patients. **e** Box plot of of the mean and 25th percentile of the expression levels (in TPM) of Basal-like non-invasive and mesenchymal-like invasive markers in basal A-like (blue) and basal B-like (red) patients. ** P <0.01, *** P <0.001, **** P <0.0001 in Wilcoxon rank-sum test comparing basal A-like to basal B-like.

When looking at the expression of well-known basal and EMT biomarkers in the two subpopulations of basal A/B-like patients, we found that basal A-like patients express classical basal/epithelial markers, such as E-cadherin, EPCAM and cytokeratins KRT5/KRT6/KRT14, together with ERBB3 and TOB1 which are markers of more differentiated, non-invasive cells (2). On the other hand, basal B-like patients express classical EMT/mesenchymal markers such as Fibronectin, the EMT inducers Twist and Slug, and the Zinc-finger transcriptional regulators Zeb1 and Zeb2 which have recently been shown to confer stemness properties that can increase the plasticity and invasive capacity of the tumour cells (50) (Fig.5d-e). In line with a more aggressive, invasive phenotype, basal B-like patients express cytoskeletal (MSN, FN1) and extracellular matrix signalling proteins (TGFB1, TGFBR2, FBN1, AXL), collagens (COL3A1, COL6A3) and proteases (MMP2, TIMP1, CTSC, PLAU, SERPINE1/2, PLAT) which are necessary for cell’s migration and invasion for dissemination to distal organs during metastasis (2). Finally, basal B-like patients overexpress a recently identified new marker of metastasis-initiating cells, the fatty acid receptor CD36 (20). Clinically, the presence of CD36 positive cells has been correlated with a lower survival rate in many carcinomas, including breast cancer, and inhibition of CD36 impairs metastasis in breast cancer-derived tumours, turning this receptor into an important biomarker of tumour cell dissemination and a potential new target to reduce cell invasion. The fact that basal B-like tumour cells co-express this metastasis-initiating marker further strengthens the aggressive nature of this tumour subclass and the clinical relevance of the basal B-specific splicing signature in tumour progression and relapse.

Overall, we have identified a novel splicing signature, specific of triple negative breast cancer tumours, that marks patients with the poorest prognosis. This basal B-like splicing signature is responsible of a stem-like, EMT phenotype that favours tumour growth, invasion of distal organs and increased drug resistance, which eventually leads to tumour relapse and metastasis. Interestingly, some of the genes differentially expressed in this basal B-like patients are well-known markers of metastasis-initiating cells, such as the alternatively spliced CTNND1 and PLOD2 genes or the fatty acid receptor CD36, turning these biomarkers into promising new targets for innovative therapies, such as the use of splicing specific antibodies (6,26).

### A metastasis-related common regulatory pathway for the basal B-specific splicing signature

Hierarchical clustering of basal A and B cell lines based on the differential expression of RNA-binding proteins highlighted six RNA regulators, ESRP1, ESRP2, RBM47, TMEM63A, KRR1 and RBMS3 (Fig. 6a) (Kruskal-Wallis p < 10^−9^).

**Figure 6.**
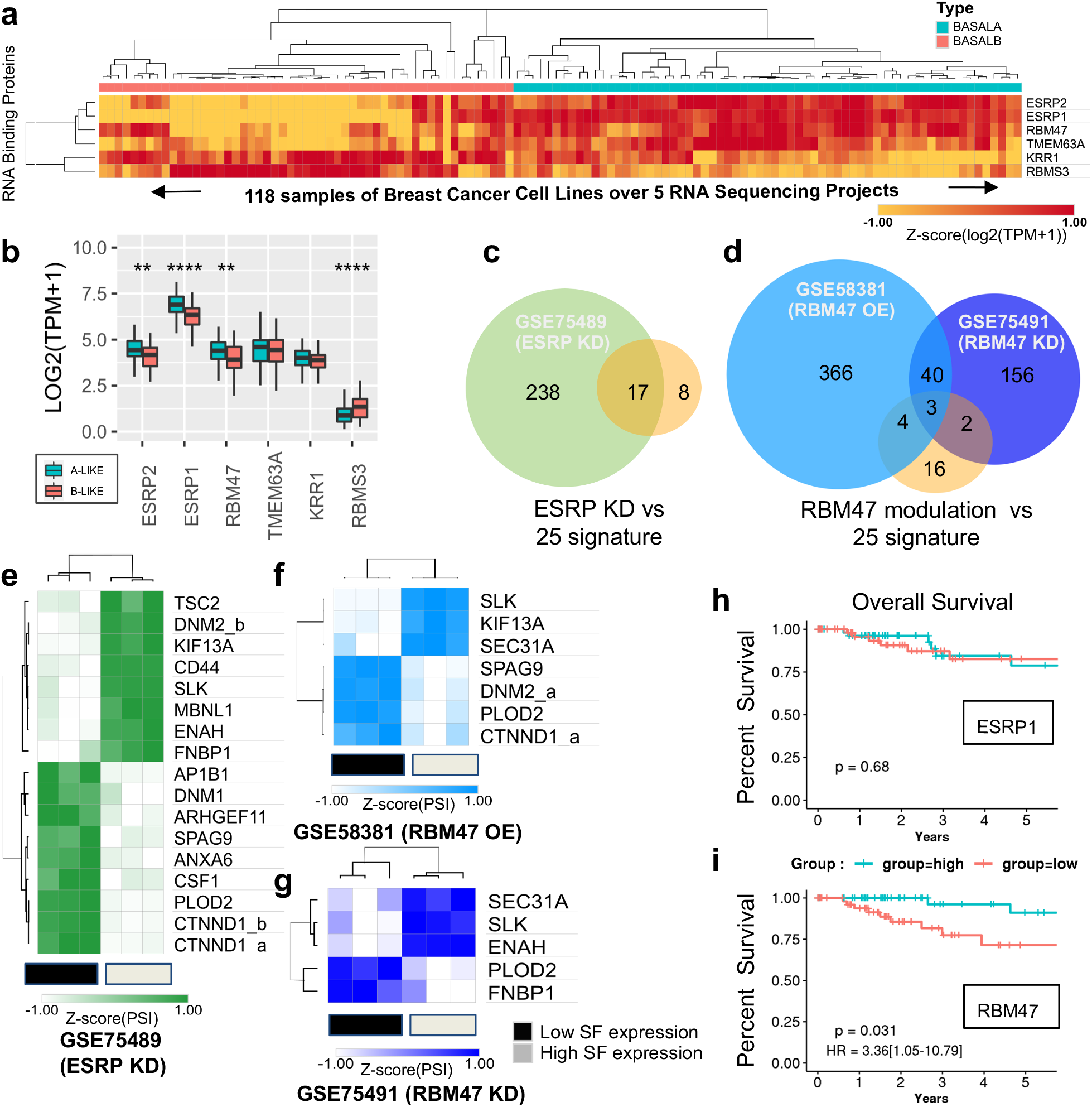
The basal B-specific splicing signature is co-regulated by ESRP1 and RBM47. **a.** Heatmap of Transcripts per Million values for RNA Binding Proteins (RBP) differentially expressed in basal A and basal B cell lines (P-value <10^−9^ by Kruskal-Wallis Test). **b**. Box plots of the mean and 25^th^ percentile of the expression levels (in TPM) of the same RBP as in **a**, but in basal A-like and basal B-like patients. **c,d**. Venn diagrams of the number of splicing events from the basal B-specific splicing signature dependent on the splicing factors (SF) ESRP1/2 and RBM47 using a cutoff of |DeltaPsi| > 0.1 and a higher probability >= 0.95. **e,f,g.** Heatmaps of the PSI values of the ESRP and RBM47-dependent exons from **c** and **d** in ESRP1/2 knock downed H358 cells, RBM47 overexpressed MDA-MB-231 cells and RBM47 knock downed H358 cells. **h,i.** Kaplan Meier plots for overall survival in basal-like TCGA patients expressing the highest tercile (blue) or the lowest tercile (red) of ESRP1 and RBM47 expression levels. HR (Hazard Ratio) and Logrank p-values (P) discriminating between groups are shown.

Interestingly, ESRP1/2 and RBM47 are significantly less expressed in basal B-like than basal A-like patients (Fig.6b), consistently with the known inhibitory effect of these three splicing regulators in EMT progression and metastasis (48,51,52). Available transcriptomics data in ESRP1/2 and RBM47 lung carcinoma NCI-H358-depleted cells (48) and RBM47 overexpressing breast cancer metastatic MDA-MB-231 cells (53) showed that 19 of the 25 splicing events responsible for the newly identified basal B-specific splicing signature are regulated by ESRP1/2 and/or RBM47 (Fig.6c-d). Importantly, ESRP1/2 and RBM47 induce the epithelial, basal A-like splicing phenotype, suggesting a potential tumour suppressor effect for these splicing regulators (Fig.6e-g, 4e and S4c-d). Consistently with this observation, low expression of RBM47 in basal-like breast cancer patients was associated with poor overall survival (log rank test p=0.031, HR=3.36, IC95%:[1.05 - 10.79] - Fig.6h-i), which supports previous experimental evidence of a role for RBM47 in supressing breast cancer metastasis and progression (52). In fact, RBM47-dependent basal B-specific splicing events were found to be functionally interconnected by physical and/or genetic interactions, which points to the existence of a common basal B-specific regulatory network associated with tumour malignancy (Fig.S5a). In support, most of RBM47-dependent basal B-specific splicing events play well-known roles in cell-cell adhesion (CTNND1) (54), cytoskeleton organization (ENAH, SLK, FNBP1) (55,56), endocytosis (KIF13A, DNM2) (57) and association with the extracellular matrix (PLOD2) (58), which are all key processes for gaining the cell motility and invasiveness necessary in tumour metastasis (54–58). Of note, expression of just one of these basal B-specific splice variants, which are CTNND1, ENAH and PLOD2, is sufficient to lower the disease-specific survival rate of basal B-like breast cancer patients compared to basal A-like (Fig.S5b-d), turning these splicing events into promising and highly specific new targets for innovative therapeutic strategies aiming at specific key splice variant instead of a pleiotropic splicing regulator that could have unsuspected secondary effects.

In summary, by taking advantage of extensive large-scale transcriptomics data from breast cancer cell lines and patients, we identified the first splicing signature capable of subclassifying basal-like tumours based on their aggressiveness and drug resistance. Importantly, novel splicing biomarkers of poor prognosis were identified that should be further studied in more functional assays to test their capacity to inhibit tumour invasion and metastasis. Results from these assays will open new perspectives in the development of improved target therapies and more accurate diagnostic profiles to identify the basal-like triple negative breast cancer patients with a higher chance of relapse.

## DISCUSSION

Cancer-specific dysregulation of alternative splicing is a promising source of cancer biomarkers and therapeutic targets to improve diagnostics and thus overall survival rate (59). An increasing number of mutations at core spliceosome components, such as S3FB1 and U2AF1, or upregulation of specific splicing factors, such as SRSF1 and other members of the SR protein family, which are now considered oncogenes, have been intimately linked to tumour progression and malignancy (60). Furthermore, an increasing number of alternatively spliced events, like CD44, ENAH, CTNND1 and FLNB, have been shown to impact cell invasion and metastasis on their own, making them promising new targets for more specific therapeutic strategies compared to the inhibition of splicing regulators (22,23,54,61,62). Effectively, splicing regulators are not only responsible for the regulation of splicing of a subset of genes, but they are also responsible for other RNA related functions such as translation, mRNA export and nonsense-mediated mRNA decay (52,60), which can have numerous downstream deleterious effects when inhibited in a targeted therapy. By specifically targeting a key downstream splicing event, as in splicing-specific immunotherapy, a more cancer-specific and direct impact on the cell phenotype might be achieved (134, 135).

Large scale public molecular data sets on genomics (copy number and mutation), epigenomics, transcriptomics, proteomics, *in vitro* and *in vivo* cell invasiveness and response to anti-tumour compounds in a large number of patients (11,000 patients across 33 different tumour types from the Genome Cancer Atlas) and human-derived cell lines (1000 cancer cell lines across 36 tumour types from the Broad Institute’s Cancer Cell Line Encyclopaedia) has become an extraordinary toolbox to identify novel prognostic markers of early metastasis and/or resistance to specific drugs, which are the two major reasons for clinical relapse and low survival rate (63–65). Unfortunately, the translatability of these pre-clinical findings is often limited since culture cells are not representative of the variety of individuals nor the biological reality of the tumour’s multicellular environment. Yet, culture procedures are improving with the creation of organoids, and machine learning approaches combined with large-scale data mining are bypassing some of these important caveats. This is the case of our cell-to-patient random forest classifier approach, in which the addition, at each round of selection, of novel informative features based on the patients classified in previous rounds allows an algorithm to make use of the information learned from cell lines. Thanks to this approach, we were able to identify the first splicing signature, composed of 25 alternatively spliced events, capable of subclassifying basal-like breast cancer patients into two subtypes with different prognoses: basal A- and basal B-like.

In fact, this newly identified basal B-like splicing signature underlined a stem-cell like EMT signature, with hallmarks of cell invasiveness and drug resistance. Five of these 25 alternatively spliced genes are well-known to play a role in cancer (ARHGEF11, CD44, CTNND1, ENAH, MBNL1) (66–68). Six have been indirectly linked to tumour malignancy and are thus new splicing targets to study (CAST, CSF1, PLOD2, SLK, SPAG9, TSC2) (56,58,69–72). The rest are completely unknown for their splicing role in cancer, even though changes in expression of some of them have been shown to play a role in tumour progression, chemosensitivity and metastasis without specifically addressing which splice variant (ATP5C1, BNIP2, FAT1, FNBP1, SEC31A, ANXA6, DNM1, DNM2) (57,73). Of special interest are ARHGEF11 and CTNND1 splice variants. Both proteins are involved in cell-cell adhesion and the basal B-specific splice variants promote cell migration and invasiveness in several cancer types, such as breast cancer (13,54,74,67). Moreover, depletion of ARHGEF11 in basal breast cancer cells is sufficient to alter cell morphology, which suppresses the cancer cell growth and survival *in vitro* and *in vivo* (67). While the existence of an isoform-specific antibody for CTNND1 pro-invasive splice variants turns this splicing candidate as a valuable new target to reduce tumour metastasis (75). ENAH and CD44 are amongst the most studied splicing events impacting cancer and are well-known biomarkers of poor prognosis. ENAH’s inhibition decreases metastasis by slowing down tumour progression and reducing cell invasion and intravasation (76–78). While the change to basal B splicing signature of CD44, a transmembrane protein that maintains tissue structure, is sufficient to drive an EMT and to increase cell invasion and plasticity by promoting stem cell characteristics (22,79). Interestingly, MBNL1 splicing regulation has also been involved in pluripotent stem cell differentiation (98–99) and cell viability via inhibition of DNA damage response (81). Promising new splice variants with a potential link with cancer are CSF1, PLOD2, SLK, SPAG9 and TSC2. CSF1 is a macrophage marker which splice variant could correlate with infiltration of tumour-promoting macrophages (69,82). Changes in the alternative splicing of the procollagen-lysine PLOD2, which catalyses the deposition and cross-link of collagens in the extracellular matrix, have been intimately linked to EMT progression and cervical, breast, lung, colon and rectal cancer prognosis (83,84). Its inhibition reduced proliferation, migration and invasion of cancer cells, while its overexpression promoted cancer stem cell properties and resistance to drugs (58,85). SLK was identified as a prognostic biomarker in several cancers and is necessary for the induction of cell migration and invasion during EMT (56,68,86). SPAG9 is a scaffold protein that organizes mitogen-activated protein kinases and has been associated with invasion in several types of tumours and prognosis (71,87,88). Finally TSC2 basal B-specific splicing isoform cannot be phosphorylated by AKT, which leads to a continuously activated mTOR pathway and oncogenic autophagy (70). More functional studies on the impact of each of these splice variants in cancer will increase our knowledge on tumour progression and metastasis with the long term goal of improving diagnostics and treatment.

It is interesting to note that these 25 alternatively spliced variants are basically dependent on three well-known splicing regulators, ESRP1/2 and RBM47, which are intimately linked to EMT and metastasis. ESRP1 is the major regulator of a newly identified epithelial-specific splicing signature (48). Its expression in cancer cells promotes tumour growth and a mesenchymal-to-epithelial transition which are essential for the formation of new tumours at distal organs during metastasis (89,90). RBM47 is a newly identified splicing regulator of EMT that has also been associated with metastasis (52,91,92). Through integrative analysis of clinical breast cancer gene expression datasets, cell line models and mutation data from cancer genome resequencing studies, RBM47 was identified as a suppressor of breast cancer progression and metastasis. It was found mutated in patients with brain metastasis and its expression was necessary to inhibit brain and lung metastatic progression *in vivo* (52). Interestingly, despite regulating just 9/25 splicing events of the basal B-specific splicing signature, low expression of RBM47, and not ESRP1, correlated with a poor prognosis and lower survival rate in basal-like breast cancer patients, which increases the interest to design new therapies targeting this splicing regulator.

In fact, this basal B-specific splicing signature has highlighted a subpopulation of basal-like triple negative breast cancer patients differentially expressing several hallmarks of invasive, EMT-like aggressive cancer, such as the newly identified biomarker of metastasis CD36 (20). CD36 is a fatty receptor expressed in metastasis-initiating cells. Neutralizing antibodies that block CD36 completely inhibited the formation of metastasis in orthotopic mouse models of human oral cancer, and CD36 inhibition impaired metastasis in human melanoma and breast cancer-derived tumours. Interestingly, the fatty acid-binding protein 7 (FABP7) correlates with a higher incidence of brain metastasis and lower survival rate in breast cancer patients, which all together points to a potential connection between fatty acid metabolism and metastasis in our subclass of basal-like breast cancer patients (93). Furthermore, cells expressing our newly identified basal B-specific splicing signature also showed resistance to several EGFR inhibiting drugs. Therapies targeting EGFR have variable and unpredictable responses in breast cancer (94). By better subclassifying sensitive from resistant tumour cells, diagnoses can be improved which will impact the choice of treatment and thus the chances of tumour relapse, or use of too severe and damaging chemotherapies. Extensive drug screening of cells derived from basal B-like patients combined with machine learning strategies to transfer the splicing knowledge will certainly improve the identification of much more suitable treatments for triple-negative breast cancer cells and reduce tumour relapse, thus improving the survival rate.

## CONCLUSION

Taking advantage of extensive available experimental data in breast cancer cell lines, we performed a knowledge transfer into clinical data to identify the first splicing signature capable of subcategorizing the most aggressive and difficult to treat type of breast cancer, which is basal-like triple negative breast cancer. Based on the pattern of splicing of 25 splicing biomarkers, we could identify two new subclasses of clinically relevant basal-like tumours, basal A and basal B-like, with different sensitivity to drugs and capacity to invade distal organs, which has a direct impact on prognosis. We propose that by testing all basal-like patients with this novel signature, patients with increased chances of creating early metastasis or tumour relapse could be closely monitored to improve their chances of survival. Similarly, by correlating alternative splicing patterns with drug resistance in cancer cell lines, or even cancer cells isolated from patients, more specific splicing biomarkers could be identified for the most adequate and personalized choice of treatment, which is one of the major challenges in triple negative breast cancer. Finally, the newly identified basal B-specific splice variants underlines a stem cell-like, highly invasive EMT phenotype, with increased drug resistance, that could be used as novel therapeutic targets to reduce cancer metastasis and relapse, opening new perspectives into the development of improved and more specific treatments for triple negative breast cancer tumours.

## METHODS

### RNA-seq transcriptomics analysis: gene expression and alternative splicing

RNA-seq reads were aligned to the human genome (GRCh38, primary assembly) using STAR (95) version 2.5.2b with standard parameters. Gencode v25 (derivated from Ensembl v85) was used for all analysis requiring annotation.

TPMCalculator (96) (v0.0.1) was used to compute Transcripts Per Million (TPM) values and obtain read counts. Q parameter was set to 255 to keep only unique mapped reads and ExonTPM value was used to consider only reads mapped to exons.

Whippet-quant from Whippet software (v10.4) was used to compute Percentage Spliced-In (PSI) values for splicing analysis. Conjointly to Kruskal-Wallis testing, the output from Whippet-quant was further filtered to include only events for which which the sum of inclusion counts (IC) and skipping counts (SC) was greater or equal to 10 for both sets of samples. Whippet-delta was used to compute differential splicing (deltaPsi) and probability that there is some change in splicing between conditions. Two heuristic filters were applied on splicing events as advised in whippet documentation; |deltaPsi| > 0.1 and P(|deltaPsi| > 0.0) >= 95 % were considered reliable parameters to filter biologically relevant AS events.

When necessary, Biobambam2 (97) (v 2.0.87) was used to transform bam files into fastq in order to be processed by Whippet.

Gene ontology (GO) analysis was done using the DAVID (v 6.8) (98) functional annotation tool (https://david.ncifcrf.gov/home.jsp) using Benjamini-Hochberg adjusted P-value cutoff of 0.05 to define a term as enriched. Go terms enrichment was restricted to GOTERM BP-FAT, GOTERM MF-FAT, and GOTERM CC-FAT, KEGG_PATHWAY and REACTOME_PATHWAY.

Gene Set Enrichment Analysis (GSEA v20.0.5) was carried out on the GenePattern (99) web platform using phenotype for permutation type and 1000 for number of permutations to execute. FDR cutoff of 25% for potential true positive finding was used as documented in the GSEA user guide. Read counts were previously normalized using DESseq2 (100) (v 1.10.1) on the same Platform.

R version 3.6.2 was used all along this study excepted for GSEA.

All heatmaps were done online using Morpheus https://software.broadinstitute.org/morpheus/. Values were adjusted by Z-score. (subtract mean and divide by standard deviation). Hierarchical clustering was done using Average linkage method and one minus pearson correlation was chosen as metric.

Sashimi plots to look cassette exons events were done using ggsashimi tool (101).

### Machine Learning and feature selection

First, we construct a classifier to distinguish basal B / A cell lines using a Random Forest with 1000 trees. After, we applied this model to the TCGA patients. Based on Gini impurity, we computed the class probability to predict patient labelled as B-like or A-like. Then, mixing initial cell lines with a subset of patients classified with the more reliability (the ones picked up with higher class probability not passing below a threshold of P=0.6), we create a new model. Each addition of patients is called a round, during which a new model is created, giving new predictions (probabilities) for the remaining patients. The algorithm stops when it can no longer incorporate the patients into one or the other group given the cut-off of P=0.6. ML analyse was done with Python 3.7.3 based on scikit-learn version 0.21.2.

To select the more efficient features that were able to separate B-like from A-like patients, we used Boruta package (0.3) implemented in python. We ran it 10 times with different random states, on the 217 features related to splicing and keep the ones that were present at least 7 times on 10. We ended with 25 AS features. Considering only these 25 AS features, we applied TSNE function from manifold package (with perplexity=20) to 3 other datasets of basal cell lines (n=56) to check the features were sufficient to distinguish spatially these cell lines according to their labels.

### Survival Analysis

Log-rank tests were performed using the functions surv and survfit from R package (survival v3.1.8). A different survival was considered significative if log rank test p-value was <0.05. Coxph function was also used for univariate Cox regression analysis in order to compute Hazard Ratio and 95% Interval of confidence. Kaplan–Meier curve were plotted using function ggsurvplot from R package survminer (0.4.6) Plots were truncated at 5 years, but the analyses were conducted using all of the data. All endpoints used for survival analysis in this study were retrieved from this study (102).

### Statistics

Wilcoxon Rank Sum Test were used to assess statistical significance within boxplots They were noted. P<0.05 (∗), P<0.01 (∗∗), and P< 0.001 (∗∗∗), P< 0.0001 (∗∗∗∗). Kruskal-Wallis Test was used to keep differential features for expression (TPM values) or splicing (PSI values) when Luminal, Basal A & B cell lines were compared and displayed in heatmap figures. A threshold of p-value <10-5 was used to filter out potential false positive and reduce the number of features in order to apply hierarchical clustering. This threshold was adapted depending on the number of samples in the comparison. For RNA binding proteins, a higher cut off of P< 10-9 was used because 5 projects were pulled together.

### Availability of data and materials

#### Raw Datasets

Five datasets of RNA-SEQ for breast cancer cell lines were downloaded. Two (PRJNA523380 and PRJNA297219) were used to transfer knowledge from cell lines to patients. Three others (PRJNA210428, PRJNA251383 and PRJEB30617) were used as a test dataset to validate importance of features. All 5 datasets were used for the RNA binding proteins analyse. The CCLE (Cancer Cell Line Encyclopedia) dataset was downloaded from the Encyclopedia GDC Legacy Archive website before availability on GEO as a BioProject (PRJNA523380). TCGA bam files for basal-like breast cancer patients were downloaded using GDC Data Transfer Tool Client from the GDC Data Portal.

Four RNA-SEQ of induced EMT were retrieved from GEO:

- HSAEC (Human Small Airway Epithelial Cells) treated with TGFB. (PRJNA260526, GSE61220)
- HMLE (immortalized human mammary epithelial cells) treated by TGFB (PRJEB25042)
- HMLE (immortalized human mammary epithelial cells) and their naturally mesenchymal CD44hi counterparts, NAMEC8 (PRJNA301721, GSE74881)
- H358 lung cells treated with doxycycline. No treatment and Day7 were kept from their timeline experiment. (PRJNA304416, GSE75492)

RNA sequencing raw reads for H358, shESRP1/2 and shRBM47 were retrieved from GEO (GSE75489 & GSE75491 respectively). RNA sequencing raw reads for MD1231-BrM2a, RBM47 induced expression were retrieved from GEO (GSE75489).

#### Breast Cancer Annotation

Basal B & A cells were labelled according to literature: Neve & al (35), Kao & al (36), Marcotte & al (103), Dai & al (104).

PAM50 intrinsic subtype were retrieved from https://www.cell.com/cancer-cell/fulltext/S1535-6108(18)30119-3 Table S4 (105).

Claudin Low status was defined with script downloaded from https://github.com/clfougner/ClaudinLow/blob/master/Code/TCGA.r (106) using dataset from http://download.cbioportal.org/brca_tcga_pan_can_atlas_2018.tar.gz (107,108).

#### Code

Code and annotation files are available here. https://github.com/LucoLab/Villemin_2020.

#### Ethics approval and consent to participate

Patients data was obtained from The Cancer Genome Atlas upon agreement of TCGA ethics and policies (https://www.cancer.gov/about-nci/organization/ccg/research/structural-genomics/tcga/history/policies)

## Availability of data and materials

RNA-seq datasets for cell lines are publicly available at https://www.ncbi.nlm.nih.gov/sra/?term=ReplaceWithID as listed in Supplementary Table S1. For patients, datasets are available at the TCGA portal upon request to https://www.ncbi.nlm.nih.gov/projects/gap/cgi-bin/study.cgi?study_id=phs000178.v11.p8

## Authors’ contributions

JPV performed all the analyses. CL helped with the development of the semi-supervised classifier. MSC and AO helped with the discussion and writing of the manuscript. JPV, RL and WR designed the study and wrote the manuscript.. All authors read and approved the final manuscript.

## Acknowledgements

We would like to thank Yaiza Nuñez-Alvarez and Sylvain Barrière for discussions.

## List of abbreviations

AS: Alternative Splicing
CE: cassette exons
EMT: Epithelial-Mesenchymal Transition
CSC: Cancer Stem Cells
CTC: Circulating Tumor Cells
PSI: Percentage Spliced-In
TPM: Transcripts per Million
DSS: Disease Specific Survival
TCGA: The Cancer Genome Atlas
RNBPs: RNA binding proteins

## Supplementary files

Supplementary Figures S1–S5

Supplementary Table S1-S2

---

**Figure S1.**
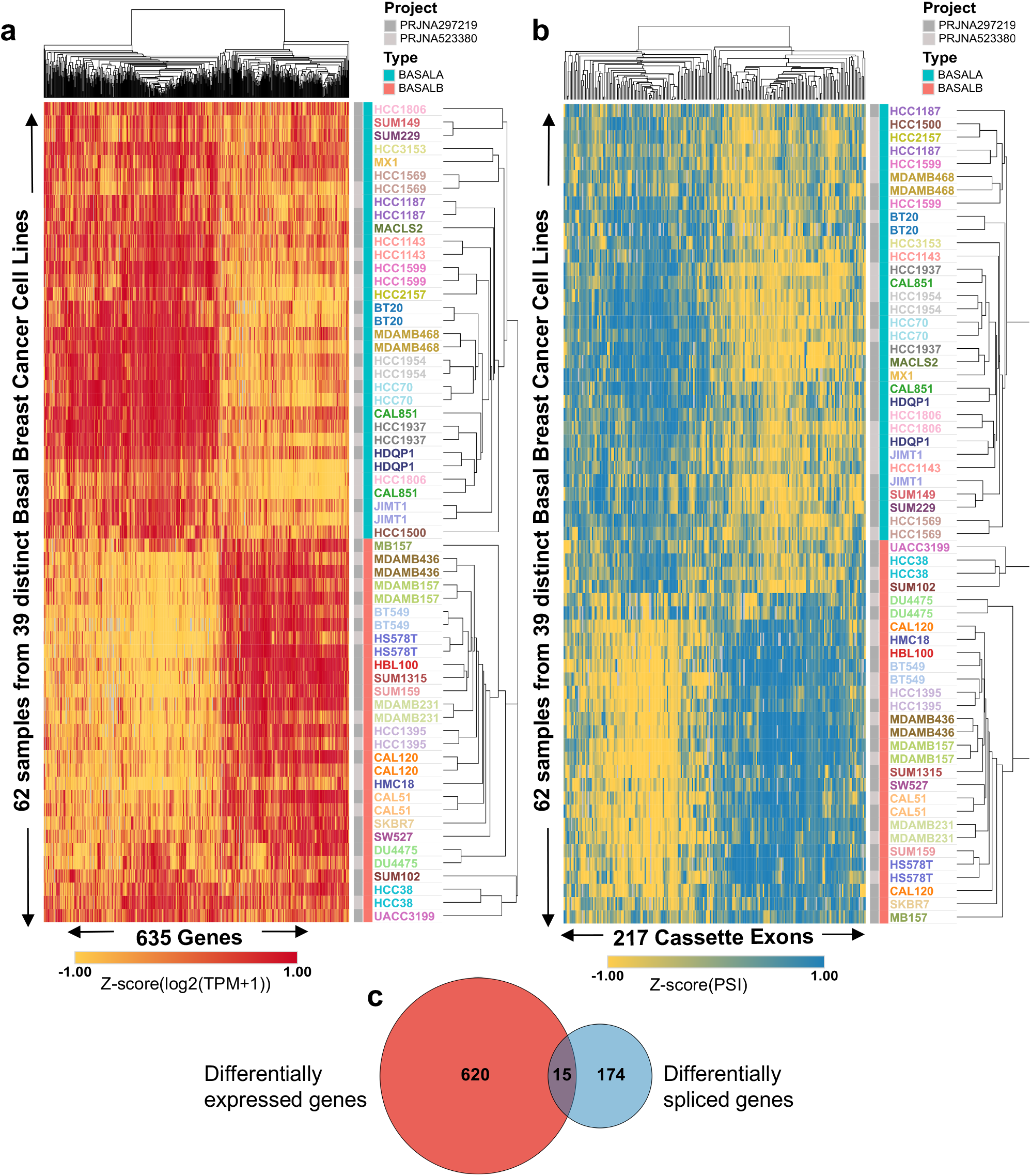
Basal cell lines are divided in two subgroups based on gene expression and splicing patterns. **a.** Heatmap of the Transcripts per Million (TPM) values of the 635 genes which differential expression can cluster breast cancer cell lines into basal A and basal B (P-value < 10^−3^ by Kruskal-Wallis Test). **b.** Heatmap of the Percentage Spliced-In (PSI) values of the 217 exons which differential splicing can cluster breast cancer cell lines into basal A and basal B (P-value <10^−3^ by Kruskal-Wallis Test). **c.** Venn Diagram of the genes differentially expressed and/or spliced between basal A and basal B cancer cell lines.

**Figure S2.**
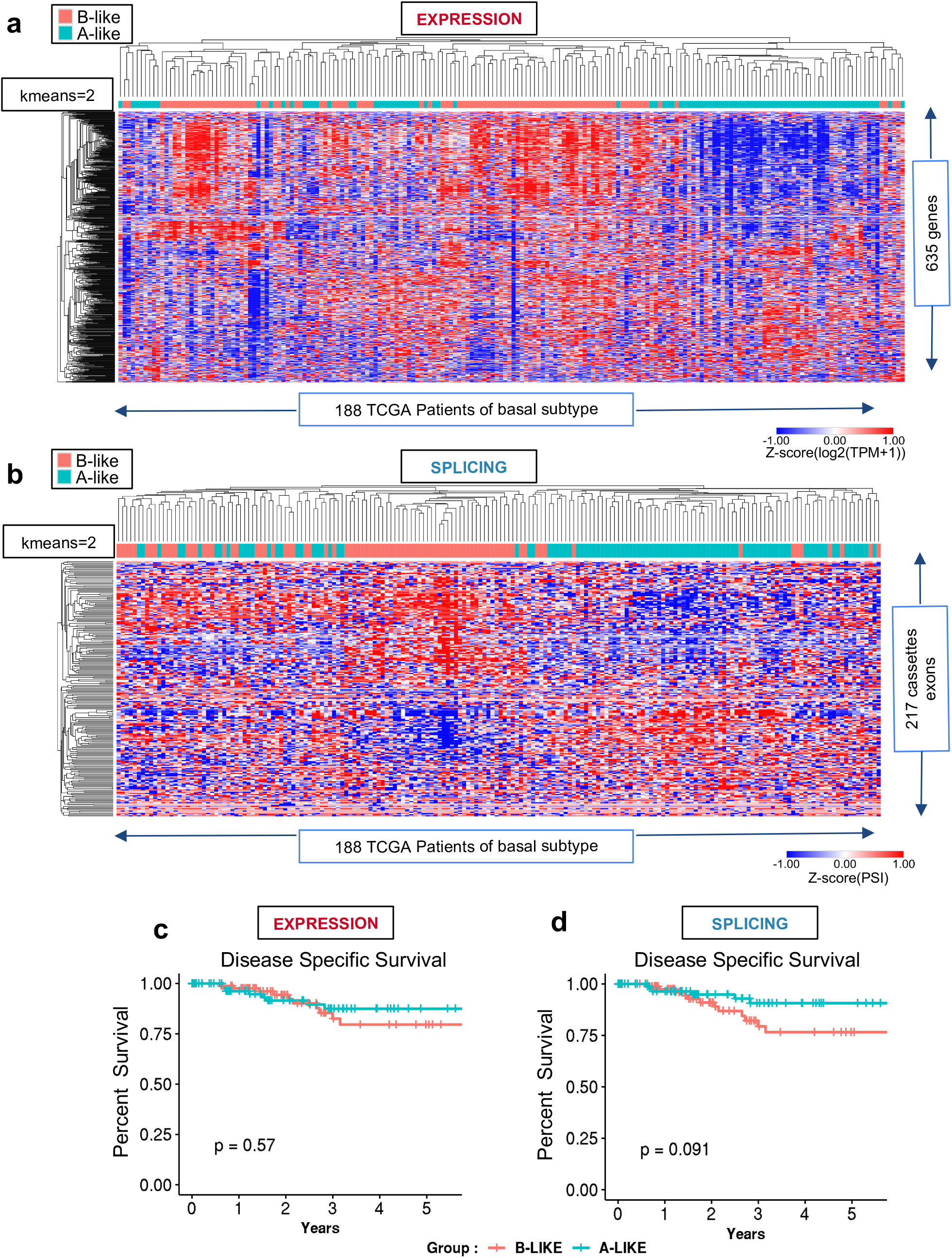
Hierarchical clustering and k-means of patients based on differential gene expression and splicing. **a-b.** Using 188 TCGA patients classified as basal-like breast cancer, we applied hierarchical clustering followed by a k-means (n=2) on expression (**a**) or splicing values (**b**) characteristic of basal B cell lines. Each time, K-means distinguished two groups we named “B-like” (red) and “A-like” (blue). In **a,** k-means was applied to TPM expression values for the 635 genes differentially expressed between basal A and B cell lines, which were displayed in the heatmap annotated Expression. In **b,** k-means was applied to PSI values of the 217 differentially spliced exons between basal A and basal B cell lines, which were displayed in the heatmap annotated Splicing. **c,d.** Kaplan-Meier plots of disease specific survival (DSS) of basal-like breast cancer patients previously separated in two groups by the k-means algorithm (k=2) for expression and splicing. Logrank test p-values (P) between “B-like” (red line) and “A-like” (blue line) patient groups are shown.

**Figure S3.**
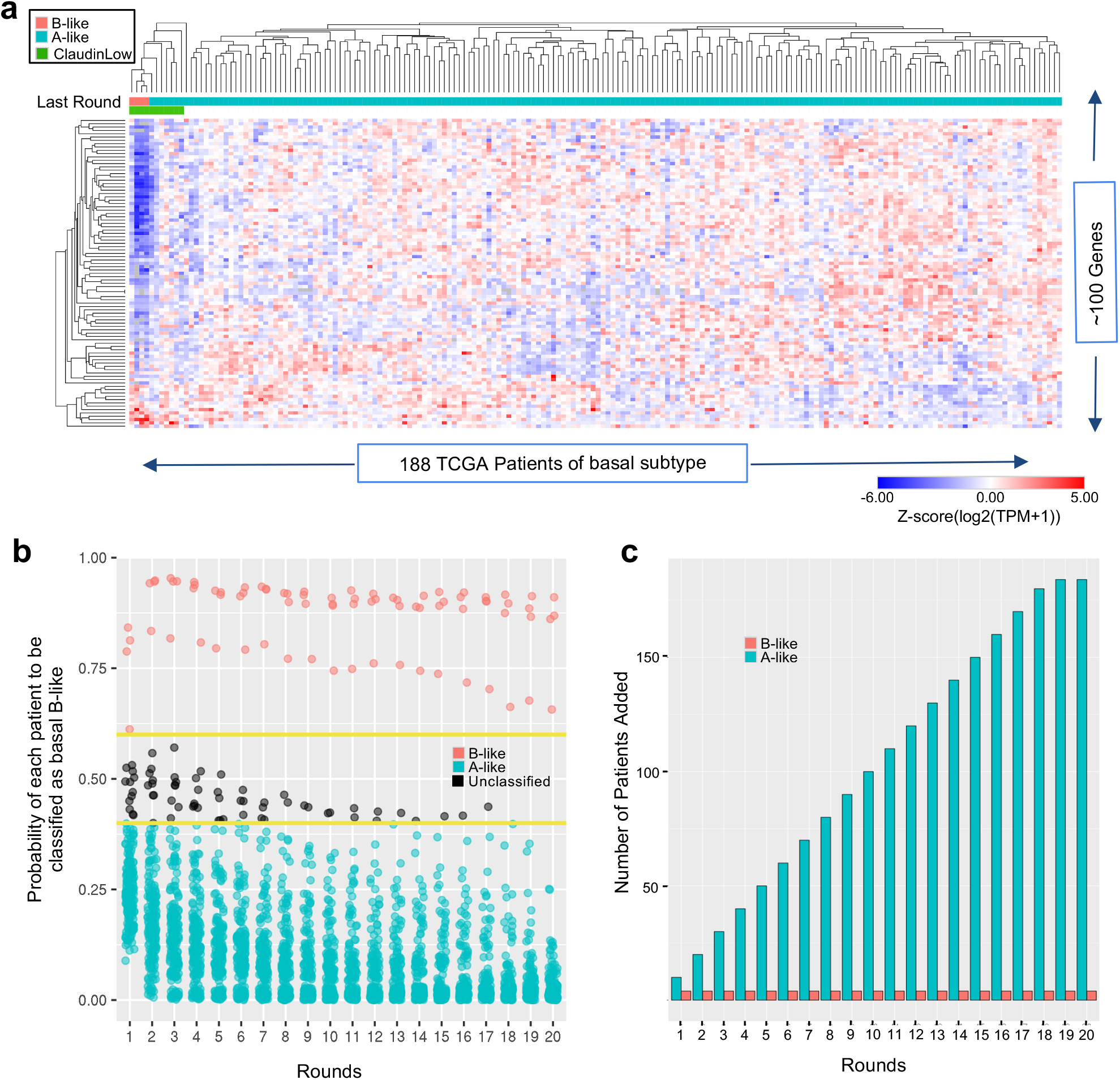
Semi-supervised Random Forest Classifier to transfer cell lines knowledge to patients using expression levels. **a.** Heatmap of ~100 genes TPM values for TCGA basal-like patients predicted as basal B-like (red) or basal A-like (blue) by the semi-supervised random forest classifier. Claudin low tumors are highlighted in green. Only the best features are represented. **b** For all patients, we plot their probabilities to be classified as basal B-like, basal A-like or unclassified at each round. Yellow lines indicate thresholds used to classify a patient as basal B-like (>0.6) or basal A-like (<0.4). **c.** Bar plot showing the number of patients added through each round. At each round, patients with the highest probability to be classified are sequentially incorporated to the initial model with cell lines in order to create a new classifier for the next round of integration.

**Figure S4.**
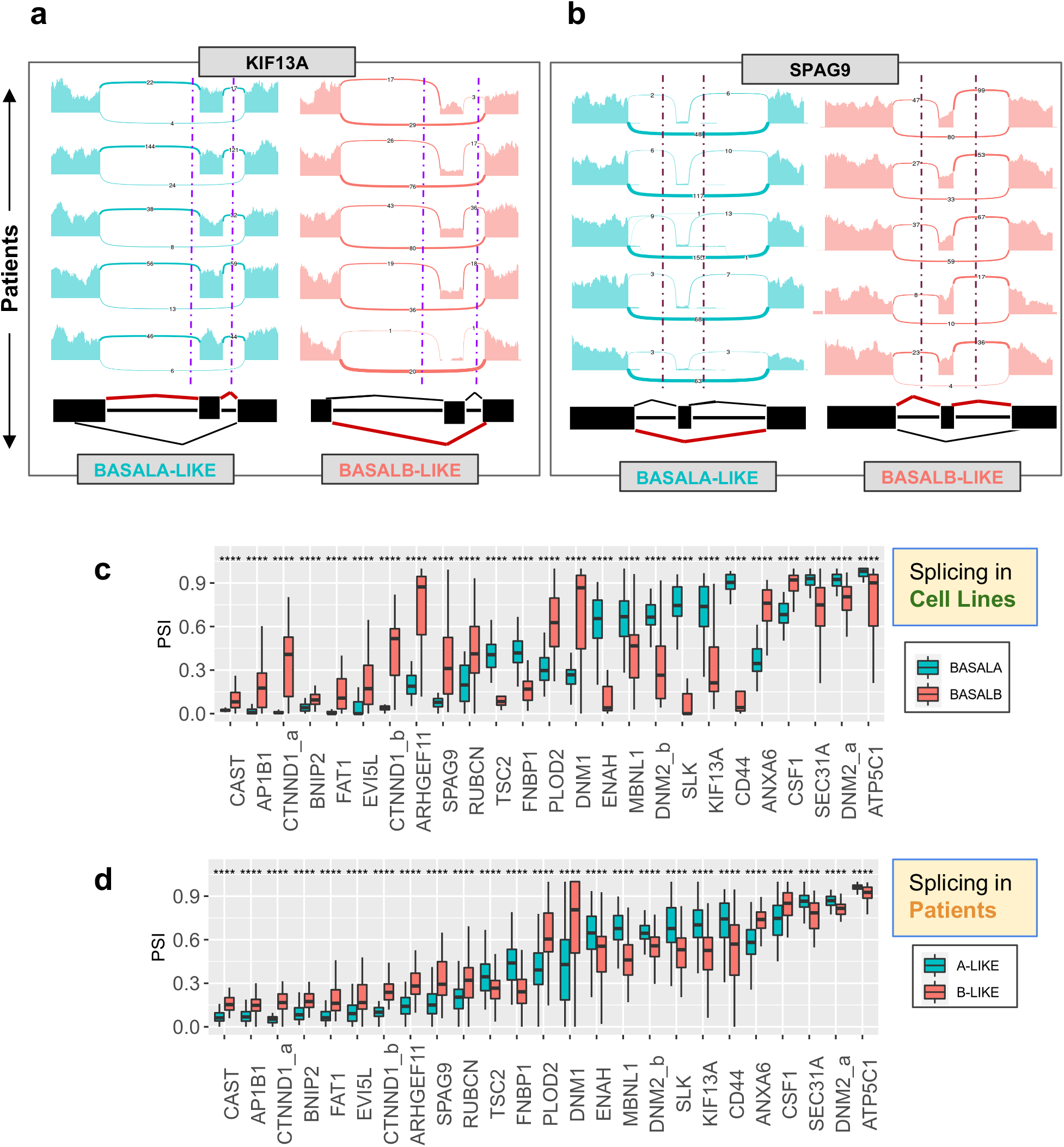
**a,b.** Sashimi plots of KIF13A and SPAG9 patterns of splicing in randomly selected basal A-like and basal B-like patients. **c,d.** Box plots of the median and 25 percentile of the Percent Spliced-In (PSI) values for the 25 cassette exons in basal A/B cell lines and basal A-like/B-like patients. **** P <0.0001 in Wilcoxon rank-sum test comparing A to B.

**Figure S5.**
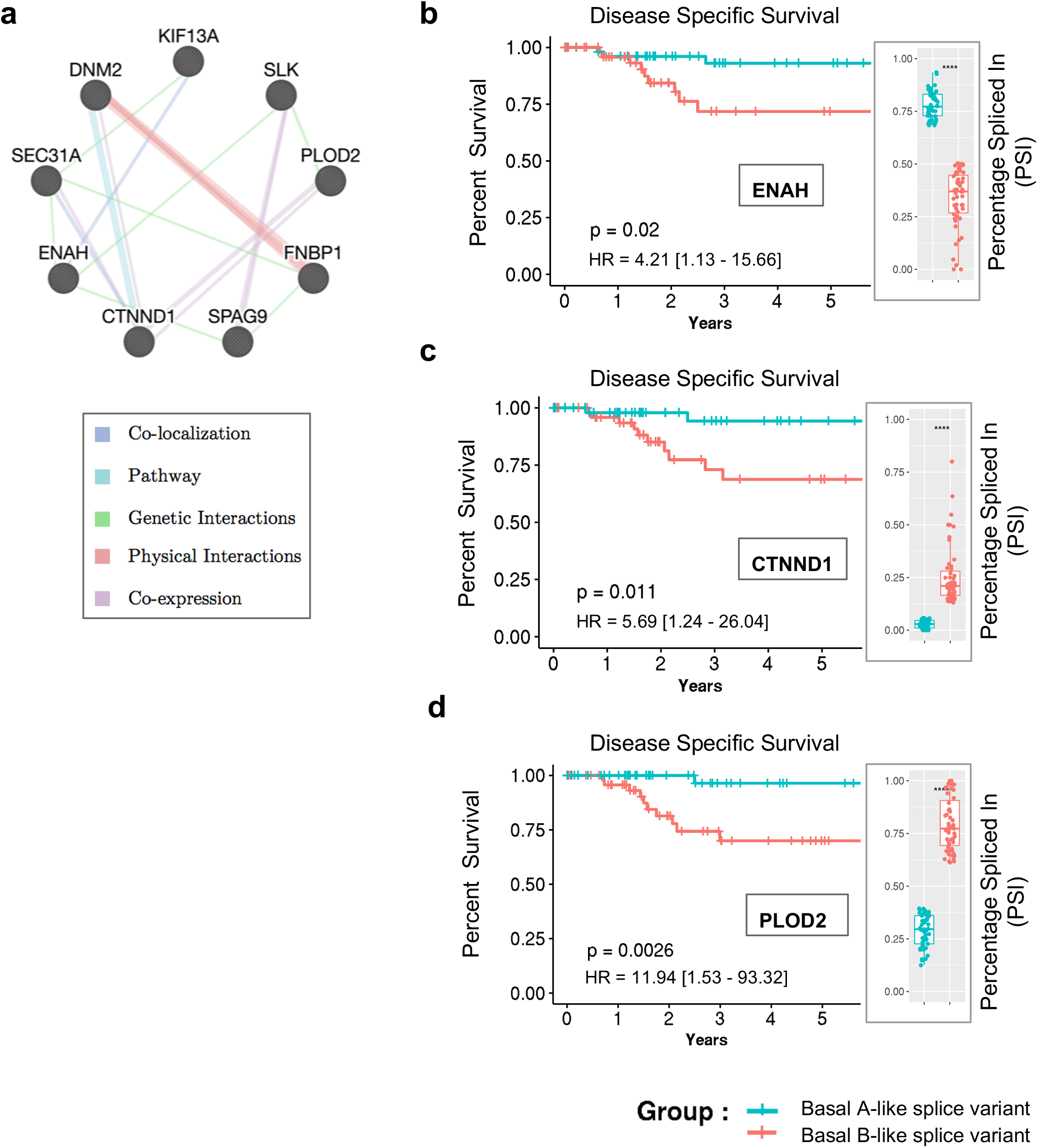
Prognostic value of individual alternatively spliced genes from the basal B-specific signature. **a**. Network of functional association (GeneMania) between RBM47-dependent spliced genes from the 25 basal B-specific splicing signature. **b,c,d.** Kaplan-Meier curves of disease specific survival in patients expressing basal A-like (blue) or basal B-like (red) ENAH, CTNND1 and PLOD2 splice variants grouped by PSI terciles. Hazard ratio (HR) and respective logrank p-values (P) discriminating groups are shown. On the left, PSI values for the first and last terciles.

